# Proteomics as a metrological tool to evaluate genome annotation accuracy following *de novo* genome assembly: a case study using the Atlantic bottlenose dolphin (*Tursiops truncatus*)

**DOI:** 10.1101/254250

**Authors:** Benjamin A. Neely, Debra L. Ellisor, W. Clay Davis

**Affiliations:** National Institute of Standards and Technology, Material Measurement Laboratory, Chemical Sciences Division, Marine Biochemical Sciences Group, Hollings Marine Laboratory, 331 Fort Johnson Road, Charleston, SC 29412, United States; National Institute of Standards and Technology, Material Measurement Laboratory, Chemical Sciences Division, Environmental Specimen Bank Group, Hollings Marine Laboratory, 331 Fort Johnson Road, Charleston, SC 29412, United States

**Keywords:** *de novo* genome, genome accuracy, proteomics, Atlantic bottlenose dolphin (*Tursiops truncatus*), marine mammal

## Abstract

**Background:** The last decade has witnessed dramatic improvements in whole-genome sequencing capabilities coupled to drastically decreased costs, leading to an inundation of high-quality *de novo* genomes. For this reason, continued development of genome quality metrics is imperative. The current study utilized the recently updated Atlantic bottlenose dolphin (*Tursiops truncatus)* genome and annotation to evaluate a proteomics-based metric of genome accuracy.

**Results:** Proteomic analysis of six tissues provided experimental confirmation of 10 402 proteins from 4 711 protein groups, almost 1/3 of the possible predicted proteins in the genome. There was an increased median molecular weight and number of identified peptides per protein using the current *T. truncatus* annotation versus the previous annotation. Identification of larger proteins with more identified peptides implied reduced database fragmentation and improved gene annotation accuracy. A metric is proposed, NP10, that attempts to capture this quality improvement. When using the new *T. truncatus* genome there was a 21 % improvement in NP10. This metric was further demonstrated by using a publicly available proteomic data set to compare human genome annotations from 2004, 2013 and 2016, which had a 33 % improvement in NP10.

**Conclusions:** These results demonstrate that new whole-genome sequencing techniques can rapidly generate high quality *de novo* genome assemblies and emphasizes the speed of advancing bioanalytical measurements in a non-model organism. Moreover, proteomics may be a useful metrological tool to benchmark genome accuracy, though there is a need for reference proteomic datasets to facilitate this utility in new *de novo* and existing genomes.

## Background

Since 2007 there has been a rapid decrease in whole-genome sequencing costs coupled with improved read lengths and development of long-range techniques such as synthetic long-reads and mapping protocols. Concurrently, the access to high performance computing environments has improved along with an endless supply of new genome assembly and annotation tools. With these new resources it is now possible to rapidly generate high-quality *de novo* genomes for non-model organisms. Excellent examples of this are two recently completed mammalian genomes (the domestic goat, *Capra hircus* [1, 2], and the Hawaiian monk seal, *Neomonachus schauinslandi* [3]) that utilized a combination of approaches including optical mapping, synthetic long reads, long read technology and chromatin interaction mapping to generate highly contiguous (scaffold N50 > 29.5 Mbp) *de novo* genomes at a relatively low cost. Overall, the result of these parallel advancements are numerous large-scale sequencing projects [4], the most ambitious targeting approximately 9 000 eukaryotic species (Earth BioGenome Project). With the forthcoming inundation of new high-quality *de novo* genomes, there is a continued need for improved metrics to evaluate genome accuracy.

Genome assemblies and annotations are evaluated in terms of contiguity and completeness, both indicators of genome accuracy. Measures of contiguity, such as scaffold N50 or N90 length, typically correspond to the quality of the genome assembly [5]. Scaffold N50 or N90 length is similar to a median or quantile scaffold length but is dependent on assembly size. Greater scaffold contiguity tends to result in more protein-coding sequences and isoforms. For example, one of the initial finished human genome assemblies from 2004 (NCBI Build 34) had a scaffold N50 of 27.2 Mbp and 27 180 protein-coding sequences, which has since been improved to a scaffold N50 of 59.4 Mbp and 109 018 protein-coding sequences (NCBI Release 108, March 2016). Gains can be even more pronounced in non-model organisms with improved *de novo* genome assemblies. For example, the *Alligator mississippiensis* (American alligator) genome recently improved from a scaffold N50 of 508 kbp to 10 Mbp using new sequencing methods [6]. Similarly, the focus of this study, *Tursiops truncatus* (Atlantic bottlenose dolphin), improved from a scaffold N50 of 116 kbp to 26.6 Mbp. Studies have shown that assembly contiguity often corresponds to assembly quality [5] but does not necessarily correlate with genome completeness and therefore accuracy [7]. One way to evaluate genome completeness is by using predicted conserved gene products. First used in the Core Eukaryotic Genes Mapping Approach (CEGMA) [8, 9], this concept has developed into Benchmarking Universal Single-Copy Orthologs (BUSCO), which is a content-based quality assessment that uses universal single-copy markers to gauge genome completeness [7]. It is evident that using many metrics to benchmark *de novo* genomes is essential to evaluating genome quality. Given the orthogonal nature of proteomics and its dependence on accurately predicted gene annotations, a quality metric based in this analytical domain may be advantageous.

Data-dependent acquisition bottom-up shotgun proteomics is one method to confirm gene annotations by observing the predicted proteins using mass spectrometry. First, proteins are digested with a known protease and the resulting peptides are fragmented within a mass spectrometer. Next, using an accurate mass of the peptide and the resulting fragmentation pattern, search algorithms can probabilistically identify peptides and then infer proteins in the search database. Alternatively, spectral libraries directly match fragmentation patterns, though these initial assignments are typically made using database-dependent approaches [10-12]. With the current generation of mass spectrometers, which have high duty cycles with high mass accuracy and resolution, we may be approaching the era of being able to infer the majority of proteins in a genome. For example, a recent proteomic analysis of HeLa tissue accounted for 91.5 % of gene products measured in the same tissue by RNA-seq (12 209 protein coding sequences versus 13 347 gene products) [13]. Since bottom-up shotgun proteomics relies completely on a database for peptide identifications and protein inference, it may be possible that a high-quality mass spectrometric dataset could be used to benchmark genome assembly and annotation quality.

The purpose of the current study was two-fold: (i) provide detailed proteomic profiling of a marine mammal and (ii) use this data to evaluate the new *T. truncatus* assembly and annotation. On average over 4 800 proteins were identified in six different tissues, and when combined yielded 10 402 protein identifications. Although not an exhaustive proteomic dataset, it confirmed approximately 1/3 of the predicted protein-coding genes. This dataset is an invaluable resource to support comparative proteomics in diving mammals related to comparative evolution [14] and biomimicry [15] and demonstrates the feasibility of accelerating cutting-edge bioanalytical approaches in non-model organisms. Secondly, the new *de novo* assembly resulted in increased protein identifications but also a decreased number of peptide identifications, despite more than a 200-fold improvement in scaffold N50 over the previous assembly. We investigated these differences at the peptide and protein level to identify global trends and proposed a new measure of genome annotation quality, NP10. This new measure was further demonstrated by evaluating human genome improvements over the past decade using publicly available proteomic data. Overall, these results highlight the improved annotation accuracy of the new *T. truncatus* genome, the utility of proteomics as a metrological tool for evaluating genome annotation quality, and emphasizes the need for reference proteomic datasets to facilitate metrology in new and existing genomes.

## Results

### Proteomic analysis of six tissues using NIST_Tur_tru v1

The initial goal of this study was to advance metrological capabilities in *T. truncatus*. This was accomplished by demonstrating proteomic measurements of six tissues from *T. truncatus*. On average, 2 199 protein groups and 4 888 proteins were identified in each tissue. The reason for performing proteomic analysis on multiple tissue types was to capture more of the possible protein population. Although there were 1 310 protein identifications shared across tissues, there was also diversity in protein identifications between tissues with the brain and skin analyses having the most unique proteins (Figure 1). Proteomic results for each tissue are available (Additional File Tables S6–S11). It is interesting to note that the liver, kidney and blubber came from the individual used for whole-genome sequencing. This dataset is relatively diverse and provides experimental evidence for over 32 000 proteotypic peptides.

**Figure 1.**
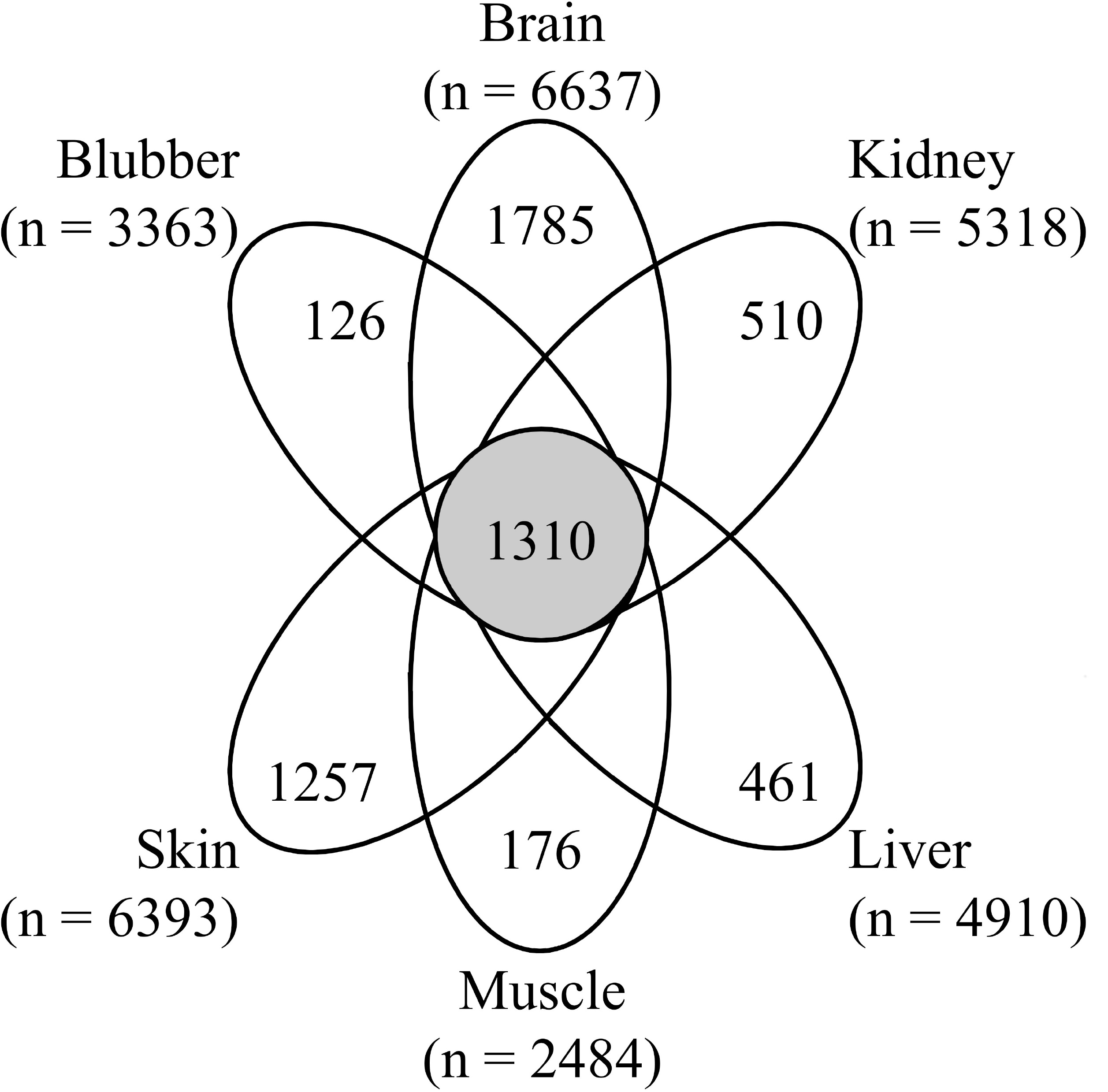
Overlap and unique protein identifications by *T. truncatus* tissue. Proteins unique to each tissue and shared by all tissues are shown along with the total number of proteins identified in each analysis.

### Comparison of Ttru_1.4 and NIST_Tur_tru v1

The second goal of the current study was to evaluate the new *T. truncatus de novo* genome assembly (GCA_001922835.1) and annotation (NIST_Tur_tru v1). This genome assembly was generated in the fall of 2016 using shotgun sequencing coupled to an *in vitro* histone ligation-based sequencing method (i.e., Chicago method) and proprietary assemblers described in detail by Putnam *et al*. [6]. This process resulted in a genome assembly with a scaffold N50 of 26.6 Mbp. Of the 159 species with genomes currently deposited on NCBI, 41 have scaffold N50 values greater than 26.6 Mbp. This level of contiguity is becoming more commonplace with three marine mammal genomes released in 2017 with scaffold N50 greater than 19 Mbp (T. *truncatus, Neomonachus schauinslandi*, Hawaiian monk seal [3], and *Delphinapterus leucas*, beluga whale [16]). For comparison, the prior NCBI *T. truncatus* annotation (Ttru_1.4) was used. This assembly was a 2012 update [14] to the 2008 draft assembly based on Sanger sequencing, Ttru_1.2 [17].

Both Ttru_1.4 and NIST_Tur_tru v1 are publicly available on NCBI and have been annotated using NCBI's eukaryotic annotation pipeline and made available in RefSeq [18]. The current annotation release, release 101 based on NIST_Tur_tru v1, has 24 026 genes and pseudogenes and 17 096 protein-coding genes with 38 849 coding sequences. At the gene and transcript level, there were many changes from Ttru_1.4 that are delineated based on alignment of genes and transcripts: identical, minor changes, major changes, new, deprecated and other. These categories are defined and available through NCBI's annotation report [19]. Briefly, 28 % of the prior genes and transcripts in Ttru_1.4 were deprecated, 72 % had minor or major changes, and 21 % of the genes and transcripts in the NIST release are new. Additionally, a small group of proteins have the prefix YP, which is not included in these NCBI categories.

Tandem mass spectrometry data collected from all six tissues was searched against each release. For both releases, almost 1/3 of the predicted protein-coding sequences were inferred by mass spectrometry. Specifically the NIST assembly identified 32 582 peptide groups belonging to 10 402 proteins comprising 4 711 protein groups. The Ttru_1.4 assembly identified 33 738 peptide groups belonging to 6 899 proteins comprising 5 292 protein groups. Many of the differences between the two results were due to a loss of deprecated sequences and minor/major changes (Figure 2). Broadly, these changes resulted in larger proteins with an increased median molecular weight and NP10 molecular weight.

**Figure 2.**
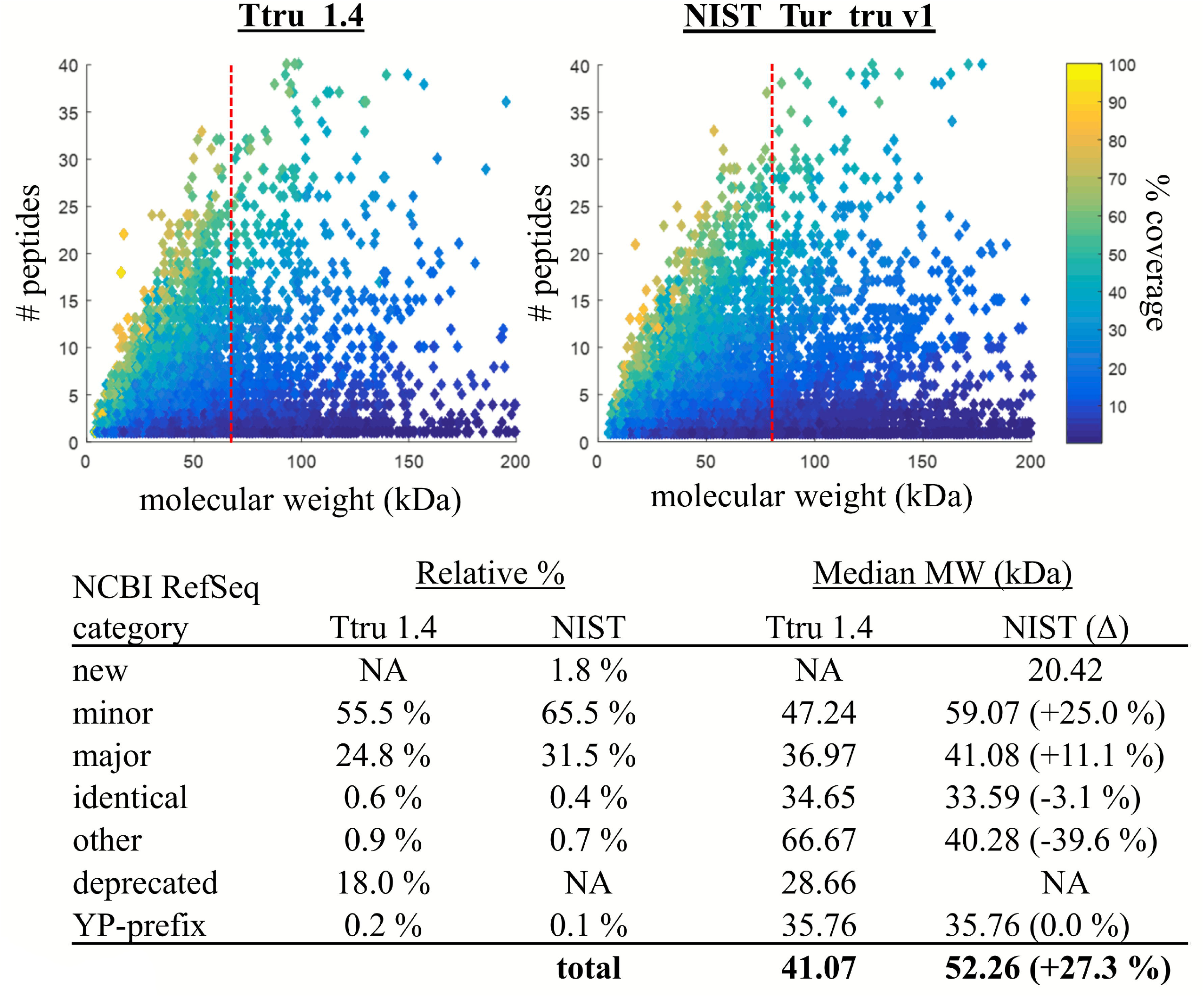
Descriptive statistics of identified proteins using different annotations. The NP10 molecular weight improved 21.3 % from 67.59 kDa to 81.99 kDa (indicated by the red dotted line) along with an improvement in median molecular weight of inferred proteins across genes with minor and major changes. (note: these axes have been truncated for illustration and do not show all data points.)

### Confirming improvements in gene annotation

There were 4,695 protein-coding sequences in the Ttru_1.4 annotation listed as partial, and one of the main improvements in the new NIST annotation was that 86 % of these sequences were merged into complete sequences. This offered an opportunity to evaluate the accuracy of these new assignments by determining whether peptides identified by mass spectrometry supported the new complete sequences. Of 6 899 identified proteins using Ttru_1.4, 1 249 were partials. Of these 1 249 partial proteins identified using Ttru_1.4, 534 had minor changes, 256 major, 450 were deprecated and 9 were other (defined simply as other changes [19]). When this NIST annotation was used, 1 005 of these same 1 249 proteins were identified, with 985 no longer being listed as partial. The median improvement within each protein was two additional unique peptides and overall the median molecular weight improved 1.8-fold (Figure 3). Of these 1 005 partial proteins identified using Ttru_1.4, when using the NIST annotation, 886 had increased molecular weight and increased number of unique peptides.

**Figure 3.**
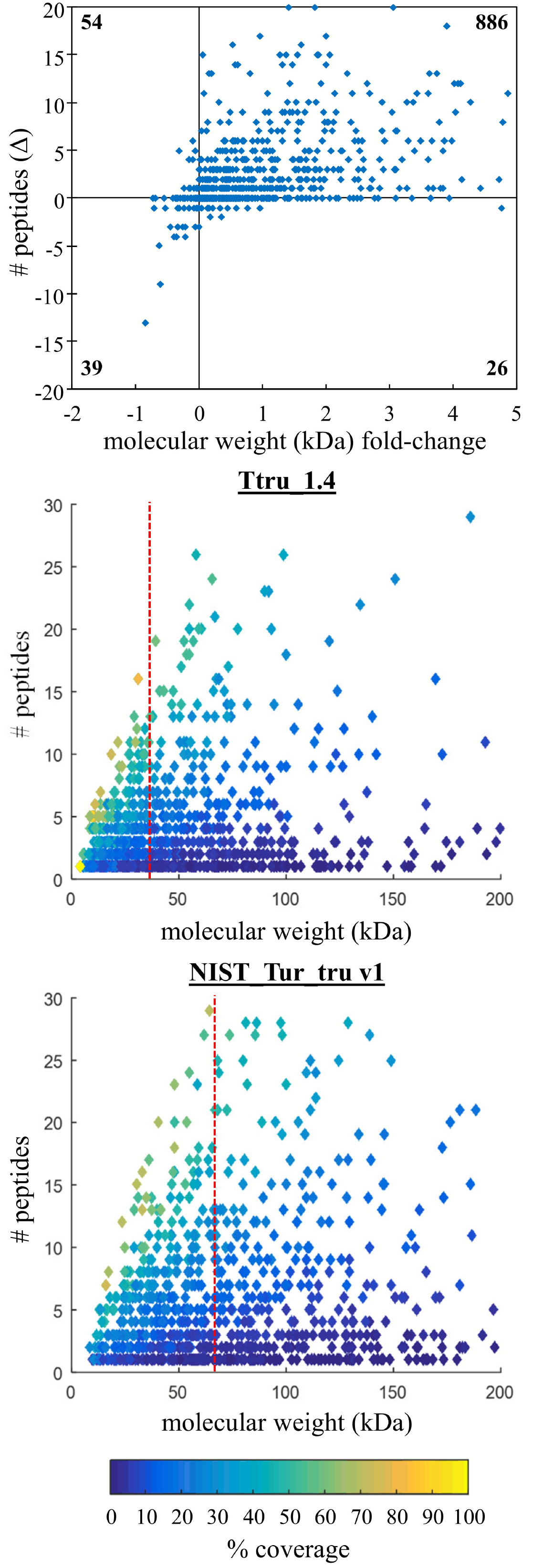
Confirming improved annotation of former partial proteins. Proteins that were partial in the Ttru_1.4 annotation were improved in the NIST annotation, and there was mass spectrometric evidence to support the accuracy of these improvements corresponding to increased peptide identifications and median molecular weight (the latter indicated by the red dotted line; note: these axes have been truncated for illustration and do not show all data points.)

### Comparing peptide identifications

An unexpected result in the new annotation was that there were fewer peptide identifications. Given the major changes between the two releases related to deprecated genes, new genes, and major changes, we were interested in tracking these peptide level changes. Over 80 % of the peptide groups identified in NIST annotation were also identified using the Ttru_1.4 annotation (Figure 4). The new peptide identifications were linked to major and minor changes in genes with only 3.2 % due to new sequences. As would be expected, many of the peptide groups not identified in the NIST annotation were deprecated (41 %). Given that these 5 657 peptide groups lost using the NIST annotation were high-confidence identifications, it may provide evidence for re-inclusion of these protein-coding sequences in future annotation releases.

**Figure 4.**
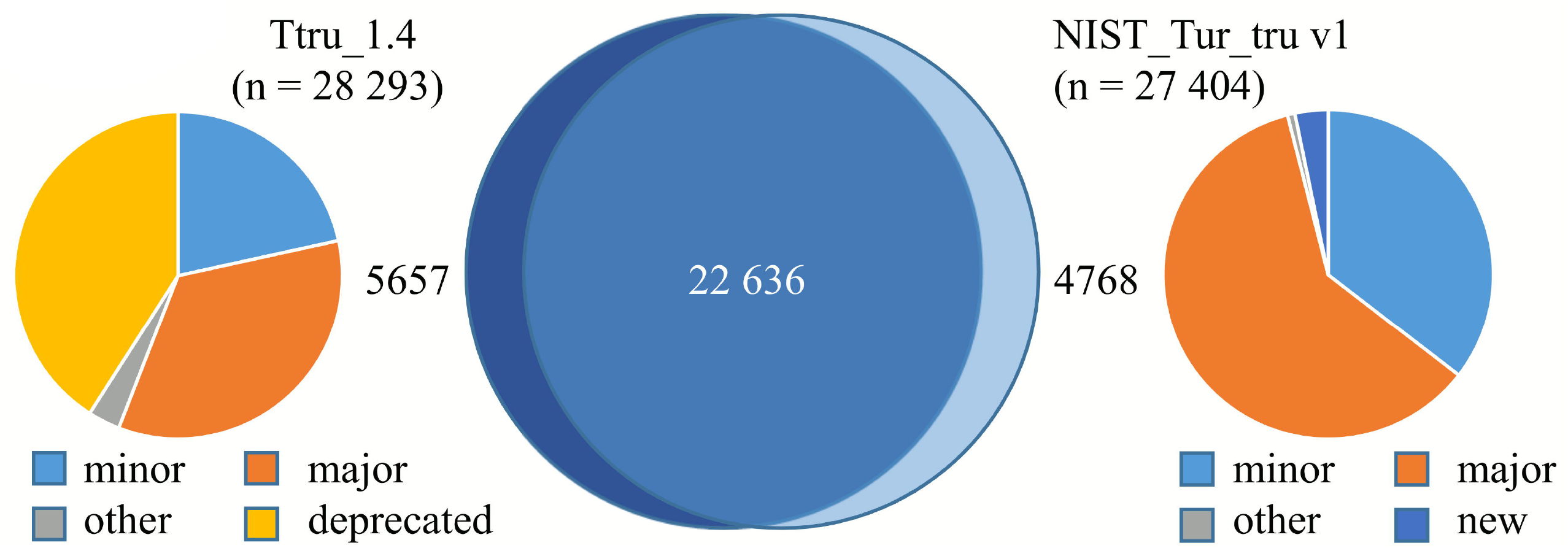
Source of peptide identification differences using the two assemblies. There was strong overlap of identified peptides using the two assemblies with over 80 % overlap. The sources of the differences were largely comprised of deprecated proteins in Ttru_1.4 (41 % of the 5 657) and minor/major changes in NIST_Tur_tru_v1 (96 % of the 4 768).

### Specific examples of annotation improvements

The goal of evaluating differences at a broad level is to capture and describe relevant changes at the granular level. At the peptide level, one the most striking improvements was related to titin, a major component in muscle tissue. In Ttru_1.4, titin (XP_004322250.1) was a partial sequence of 2,167 amino acids (241.7 kDa) and 60 unique peptides (40.2 %) were identified belonging to this sequence. In the NIST annotation, the coding sequence for titin (XP_019787158.1) was 32 192 amino acids (3 812.8 kDa) and 779 unique peptides (34.3 % coverage) were identified belonging to this sequence. This single sequence improvement is responsible for many changes observed at the peptide level (Figure 4).

Almost 2 % of the identified proteins using the NIST annotation were considered new. One important new protein of note is cystatin C (XP_019783122.1). This protein was not present in Ttru_1.4, while using the NIST annotation the mass spectrometry data identified three unique peptides (41.3 % coverage) belonging to the predicted 13.1 kDa protein. This protein has applications as a biomarker [20], and with these proteomic results, it is possible to create SI traceable mass spectrometer-based assays (similar to [21]). Another protein of note is serotransferrin (XP_019789750.1), which is 90 % identical and 3.5 % longer than the entry in Ttru_1.4 annotation (XP_004329553.1). Most of these changes were on the c-terminus section (from positions 537 to 634), which was supported by the proteomic data that identified four peptides spanning this region. There were other slight changes to the sequence that resulted in six more unique peptides identified in the improved serotransferrin, which supports the accuracy of the new annotation. Overall, there are many changes related to the over 10 000 protein identifications and many would be considered improvements as indicated by increased protein molecular weight and/or greater peptide coverage. At a gene-by-gene level these results can be used to confirm and improve annotations.

### Confirming quality metric in human annotations

In order to gauge the broader applicability of using proteomics as a quality measure of genomic annotations, we demonstrated NP10 in a more mature genome with deeper proteomics. The recent work by Bekker-Jensen *et al*. [13] is publicly available on ProteomeXchange [22, 23] and for this comparison the data generated from a 39 fraction high pH pre-fractionation of a HeLa cell digest followed by LC-MS/MS analysis was used for database searching. These data were searched against three human genome annotations from 2004, 2013 and 2016, each with markedly increased scaffold N50 values and database sizes (i.e., number of coding-sequences; Table 1). The number of identified proteins was 13 341, 22 906, and 48 019 proteins in Build 34, Release 105 and Release 108, respectively. The median molecular weight improved 25 % (from 51.06 to 53.46 to 63.99 kDa, respectively) whereas the improvement in NP10 was more pronounced with a 33 % improvement (from 100.17 to 101.87 to 133.55 kDa, respectively; Figure 5).

**Table 1.** Descriptive statistics of human annotated databases and resulting proteomic identifications.

|  | Build 34 | Release 105 | Release 108 |
| --- | --- | --- | --- |
| **release date** | Feb 2004 | Jun 2013 | Mar 2016 |
| **scaffold N50** | 29.1 Mbp | 45.0 Mbp | 59.4 Mbp |
| **coding sequences** | 27 180 | 45 107 | 109 018 |
| **protein groups** | 9 762 | 10 059 | 10 219 |
| **proteins** | 13 341 | 22 906 | 48 019 |
| **peptide groups** | 175 895 | 184 580 | 184 806 |
| **peptide spectral matches** | 390 909 | 405 852 | 405 950 |

**Figure 5.**
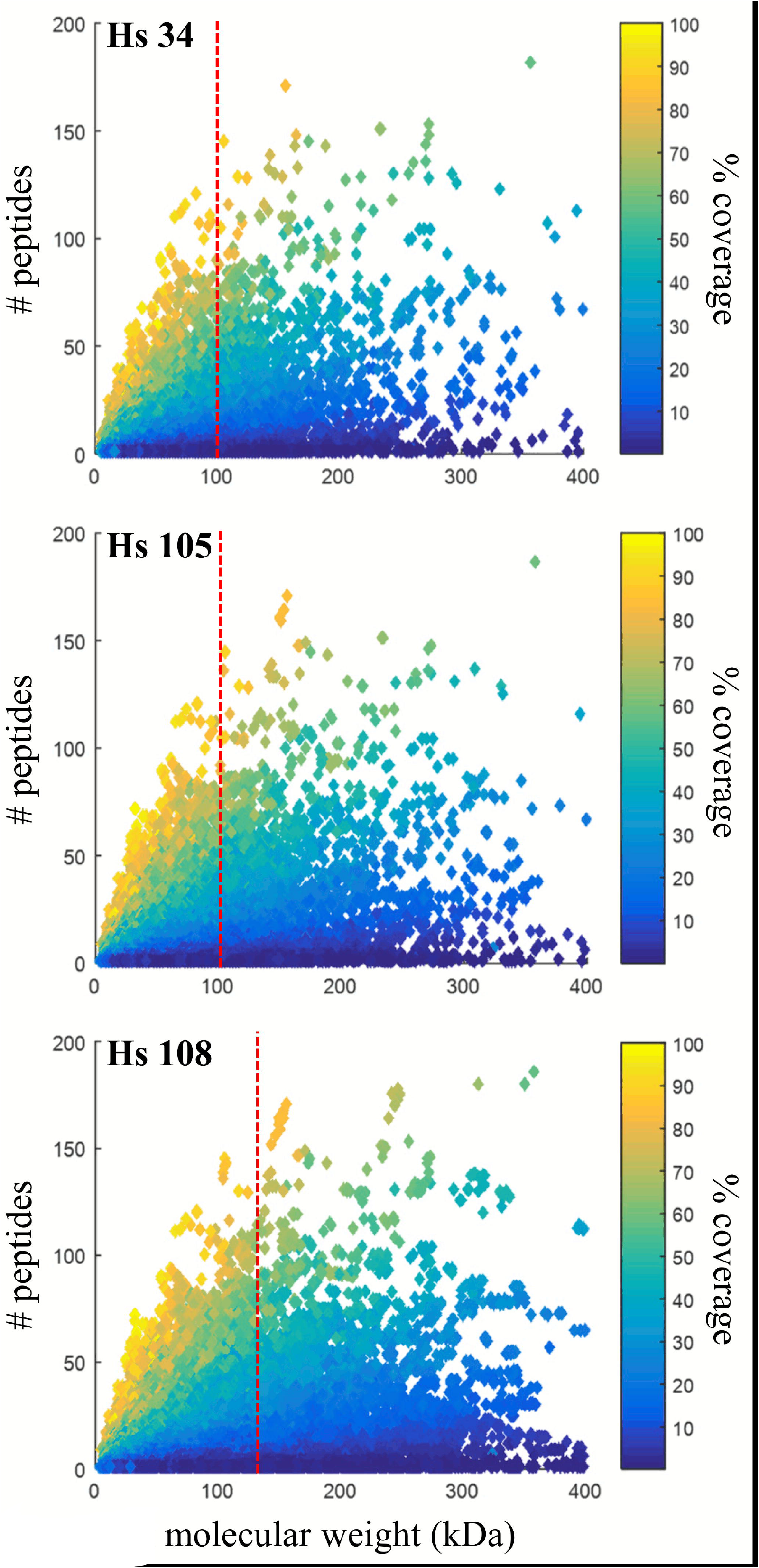
Similar trends with improved human assemblies. As the contiguity of the human genome has improved, there is a shift upward and to the right indicating annotations are more accurate (increased coverage) and complete (increased molecular weight). The NP10 improved 33 % and is indicated by the red dotted line (note: these axes have been truncated for illustration and do not show all data points).

## Discussion

Advances in bioanalytical platforms across domains (*i.e*., genomics, transcriptomics, and proteomics) are improving the accessibility of non-model organisms as viable research candidates. The results of the current study provide secondary confirmation of 10 402 proteins from 4 711 protein groups using a recently completed well-scaffolded high-coverage *T. truncatus* genome and shotgun proteomic analysis of six different tissues. Previous proteomic studies of *T. truncatus* have identified less than 100 protein groups in serum [15, 21], while the most detailed published proteomic analysis of a marine mammal identified 206 proteins in cerebrospinal fluid of *Zalophus californianus* (California sea lion) [24]. Currently there are twelve marine mammal genomes that have been annotated by NCBI (of the 159 species with genomes currently deposited on NCBI), though only *T. truncatus* and *Z. californianus* have published mass spectrometry based proteomic datasets. Work is underway to increase the number of marine mammal genomes along with companion high-quality proteomic datasets and spectral libraries. The results of the current study provide empirical confirmation of protein annotations, including observable proteotypic peptides, which can be a resource for future targeted studies in *T. truncatus*. For example, by improving the protein-coding sequence accuracy of serotransferrin in *T. truncatus*, future studies can extrapolate metrological advances in human serotransferrin sialoforms [25] to *T. truncatus* disease treatment [26]. Since the current results are not an exhaustive proteomic dataset, future studies will utilize different solubilization techniques, proteases, and separation techniques to provide even deeper proteome coverage (reviewed and demonstrated in the following [13, 27, 28]). Still, it is worth noting that in single study using a simple experimental approach we have identified almost 1/3 of the possible predicted proteins, emphasizing the ease of accomplishing bioanalytical advances in non-model organisms using modern techniques.

In the current study, benchmark proteomic datasets were used to evaluate genome assembly and annotation improvements in *T. truncatus* and *H. sapiens*. Typically, a reference database is used to demonstrate proteomic improvements due to optimized protein extraction, solubilization and digestion, peptide separation, mass spectrometer speed and mass accuracy, search algorithm performance and database accuracy. In contrast, when the mass spectrometric data are held constant and instead the database is varied, differences in proteomic results are indicative of database fragmentation and accuracy. Proteomic analysis of multiple tissues allowed for greater protein diversity when evaluating *T. truncatus*, though the publicly available human data performed exceptionally well despite using a single tissue since it utilized highly optimized separation techniques. An optimum proteomic benchmark dataset would be one that offers the possibility of the deepest proteome coverage. This would rely on using multiple tissues, extraction protocols, enzymes and optimum separation techniques coupled to modern mass spectrometers. These datasets could be developed in parallel to the exponential increase in *de novo* genomes being released and annotated and would prove invaluable in exercises assessing assembly and annotation performance (such as Assemblathon 2 [5]). Importantly, given the abundance and accessibility of public proteomic data in this “Golden Age of Proteomics” (as coined by [29]) and modular open-access proteogenomic pipelines such as Galaxy-P [30, 31], it would be possible to incorporate these reference mass spectrometric datasets and proteomic derived quality metrics into genome assembly and annotation pipelines.

In parallel to improvements in genome assembly contiguity and annotation accuracy, proteomic results should have increased peptide numbers per protein, higher protein identifications due to isoform resolution and improved coverage of higher molecular weight proteins due to better long-range accuracy. For instance, when evaluating the substantial reduction in partial sequences between Ttru_1.4 and NIST_Tur_tru v1, there was an increase of 81 % in median molecular weight of these proteins that coincided with more peptide identifications within these new complete sequences. The most drastic example in this case study was titin, which went from 60 to 779 identified peptides with the addition of over 32 000 amino acids to the previously partial sequence. This also emphasizes that greater numbers of protein identifications does not imply higher quality since a more fragmented genome will give more protein identifications. Instead, identification of larger proteins with more identified peptides is more indicative of improved quality. The proposed metric, NP10, attempts to capture this quality measure. One issue is that the NP10 may be glossing over how changes in spectral assignments to peptides with changing databases affect proteomic quality (such as false discovery rates). There is an opportunity to develop a streamlined method to track MS/MS spectra assignments and quantify those changes with database improvements in order to establish finer measures of search space effects on proteomic performance. Overall, these results demonstrate that new whole-genome sequencing techniques can provide high quality *de novo* genome assemblies and that proteomics is a useful metrological tool to evaluate annotation and benchmark genome accuracy.

## Methods

### Sample source and preparation

Bottlenose dolphin tissues were collected from animals under appropriate permits (Additional File Table S1) and stored at liquid nitrogen temperatures (−150 to −180 °C) until cryohomogenization in the National Institute of Science and Technology's Marine Environmental Specimen Bank [32]. From the resulting fine powder, 5 mg was subsampled and the proteins were extracted using RapiGest (Waters, Milford MA). Briefly, 150 μL of 0.1 % (w/v) RapiGest (in 50 mM ammonium bicarbonate) was added, resulting in a solution of 33 μg/μL tissue. The extraction mixture was shaken at 600 rpm for 25 min at room temperature followed by removal of large debris using a benchtop microcentrifuge. From this solution, a 5 μL aliquot was removed and suspended in 35 μL of 0.1 % (w/v) RapiGest (in 50 mM ammonium bicarbonate), followed by the addition of 40 uL of 50 mM ammonium bicarbonate. Next, the sample was reduced with 10 μL of 45 mM dithiothreitol (DTT; final concentration of 5 mM) and incubated at 60 °C for 30 min, then allowed to cool to room temperature. The mixture was alkylated using 3.75 μL of 375 mM iodoacetamide (Pierce, Thermo Scientific, Waltham, MA; final concentration of 15 mM) and incubated in the dark at room temperature for 20 min. Prior to addition of trypsin, 100 μL of 50 mM ammonium bicarbonate was added. A 3.3 μL aliquot of trypsin (MS-Grade; 1 μg/μl in 50 mM acetic acid) was added (1:50 trypsin:protein) and samples were incubated overnight at 37 °C. The digestion was halted and RapiGest cleaved with the addition of 100 μL of 3 % (v/v) trifluoroacetic acid (1% final concentration) and incubated at 37 °C for 30 min before centrifiigation and removal of the supernatant. Samples were processed using Pierce C18 spin columns (8 mg of C18 resin; Thermo Scientific) according to manufacturer's instructions. Each sample was processed in duplicate yielding at maximum of 60 μg peptides. These solutions were evaporated to dryness in a vacufuge then reconstituted in 150 μL of 5 % acetonitrile in water.

### Mass Spectrometry

Samples were analyzed using an UltiMate 3000 Nano LC coupled to a Fusion Lumos mass spectrometer (Thermo Fisher Scientific). Resulting peptide mixtures (10 μl) were loaded onto a PepMap 100 C18 trap column (75 μm id x 2 cm length; Thermo Fisher Scientific) at 3 μL/min for 10 min with 2 % (v/v) acetonitrile and 0.05 % (v/v) trifluoroacetic acid followed by separation on an Acclaim PepMap RSLC 2 μm C18 column (75 μm id x 25 cm length; Thermo Fisher Scientific) at 40 °C. Peptides were separated along a 130 min gradient of 5 % to 27.5 % mobile phase B [80 % (v/v) acetonitrile, 0.08 % (v/v) formic acid] over 105 min followed by a ramp to 40 % mobile phase B over 15 min and lastly to 95 % mobile phase B over 10 min at a flow rate of 300 nL/min. The mass spectrometer was operated in positive polarity and data dependent mode (topN, 3 s cycle time) with a dynamic exclusion of 60 s (with 10 ppm error). The RF lens was set at 30 %. Full scan resolution using the orbitrap was set at 120 000 and the mass range was set to *m/z* 375 to1500. Full scan ion target value was 4.0e5 allowing a maximum injection time of 50 ms. Monoisotopic peak determination was used, specifying peptides and an intensity threshold of 1.0e4 was used for precursor selection. Data-dependent fragmentation was performed using higher-energy collisional dissociation (HCD) at a normalized collision energy of 32 with quadrupole isolation at *m/z* 0.7 width. The fragment scan resolution using the orbitrap was set at 30 000, *m/z* 110 as the first mass, ion target value of 2.0e5 and a 60 ms maximum injection time.

### Protein Search parameters

Resulting raw files from the analysis of six different *T. truncatus* tissues and raw files from a publicly available 39 fraction HeLa experiment (ProteomeXchange Consortium [23] via the PRIDE partner repository with the dataset identifier PXD004452) were processed and searched using Proteome Discoverer (v.2.0.0.802). For *T. truncatus* analysis, Sequest HT and Mascot (v2.6.0; Matrix Science) search algorithms were used, while only Sequest HT was used for human searches. For all searches, the protein.faa fasta file was retrieved from NCBI RefSeq [18] via ftp [33]. For searches with the prior *T. truncatus* annotation, GCF_000151865.2_Ttru_1.4 was used, while searches with the current *T. truncatus* annotation, GCF_001922835.1_NIST_Tur_tru_v1 was used. These correspond to release 100 and 101 for this organism on NCBI. The whole-genome sequencing projects can be found in GenBank [34] under entries ABRN00000000.2 (Ttru_1.4) and MRVK00000000.1 (NIST_Tur_tru_v1). For the human searches, the following were used: GCF_000001405.10_hg16_Build34.3 (Build 34), GCF_000001405.25_GRCh37.p13 (Release 105) and GCF_000001405.33_GRCh38.p7 (Release 108). The *T. truncatus* searches also used the common Repository of Adventitious Proteins database (cRAP; 2012.01.01; the Global Proteome Machine), though these sequences were removed from search results.

The following search parameters were used for Mascot and Sequest: trypsin was specified as the enzyme allowing for two mis-cleavages; carbamidomethyl (C) was fixed and acetylation (protein n-term), deamidated (NQ), pyro-Glu (n-term Q), and oxidation (M) were variable modifications; 10 ppm precursor mass tolerance and 0.02 Da fragment ion tolerance. Within Sequest, the peptide length was specified as a minimum of six and maximum of 144 amino acids. Resulting peptide spectral matches were validated using the percolator algorithm, based on q-values at a 1 % false discovery rate (FDR). The peptides that were greater than six amino acids long were grouped into proteins according to the law of parsimony and filtered to 1 % FDR and single peptide hits were allowed. Briefly, there may be more than one peptide spectral match for a given peptide, which are then grouped to peptide groups. Protein inference is when these peptide groups are assigned to proteins, but given similarity between some proteins (such as isoforms or highly homologous sequences), peptides can match to more than one protein. For this reason, protein families or protein groups are generated based on peptide overlap (and therefore sequence overlap), which reduces inflation due to isoform identifications. For the described analyses, protein and peptide groups are used and are available for each *T. truncatus* search in Additional File Tables S2 – S5. Raw MS data and Mascot based search results for *T. truncatus*, as well as all fasta databases, have been deposited to the ProteomeXchange Consortium [23] via the PRIDE partner repository with the dataset identifier PXD008808 and 10.6019/PXD008808.

### Proteomic-based quality metric for annotation quality

Evaluating proteomic results relies on qualifying how well a database explains the observed tandem mass spectra: high numbers of protein identifications and percent identified spectra indicate good proteomic performance. Another way of describing proteomic results is to plot the number of peptide identifications versus protein molecular weight. A larger protein has potentially more peptide identifications but due to solubilization and digestion effects (such as post-translational modifications and protein folding), larger proteins do not always yield more unique peptides. For this reason, there is a somewhat Gaussian distribution of peptide frequency around median protein molecular weight. This median can shift right when the molecular weight of predicted protein-coding sequences increases and/or the number of isoforms increases.

When evaluating and comparing *de novo* genome assemblies and annotations, the specific question that proteomics can answer is the degree of database fragmentation and accuracy. If an annotation improves partial coding sequences to complete protein-coding sequences with isoforms, then there will be an increase in the molecular weight of identified proteins with more peptides assigned to these longer sequences. By simply improving partial sequences there would be a shift to higher protein molecular weight. One goal of the current study was to provide a more robust quality measure by incorporating unique peptide counts (which corresponds to protein coverage) with the change of median molecular weight of inferred proteins. The NP10 is a proposed metric that first stratifies the results by identifying the top decile (or 10^th^ 10-quantile) of proteins based on the number of peptides per protein and then returns the median molecular weight of the resulting proteins (graphically demonstrated in Additional File Figure S1). This metric is similar to simply calculating the median molecular weight of all inferred proteins, but by removing protein identifications with relatively few peptide assignments, it attempts to indicate accuracy of the improved/longer protein-coding sequences.

## Availability of supporting data

The raw data and tissue specific search results along with all databases used are available at the ProteomeXchange Consortium [23] via the PRIDE partner repository with the dataset identifier PXD008808 and 10.6019/PXD008808. The proteomic data from Bekker-Jensen *et al*. [13] used for the human comparison can be found at ProteomeXchange Consortium [23] via the PRIDE partner repository with the dataset identifier PXD004452. Tabulated search results for combined analysis and for each tissue can be found in Additional File Supplemental Tables S1-S11.

> Additional File Figure S1. Graphical example of NP10 calculation.
>
> Additional File Table S1. Sample characteristics table.
>
> Additional File Table S2. Protein Identifications using Ttru_1.4.
>
> Additional File Table S3. Protein Identifications using NIST_Tur_tru v1.
>
> Additional File Table S4. Peptide Group Identifications using Ttru_1.4.
>
> Additional File Table S5. Peptide Group Identifications using NIST_Tur_tru v1.
>
> Additional File Table S6. Protein Identifications in blubber tissue using NIST_Tur_tru v1.
>
> Additional File Table S7. Protein Identifications in brain tissue using NIST_Tur_tru v1.
>
> Additional File Table S8. Protein Identifications in kidney tissue using NIST_Tur_tru v1.
>
> Additional File Table S9. Protein Identifications in liver tissue using NIST_Tur_tru v1.
>
> Additional File Table S10. Protein Identifications in muscle tissue using NIST_Tur_tru v1.
>
> Additional File Table S11. Protein Identifications in skin tissue using NIST_Tur_tru v1.

## Declarations

### List of abbreviations

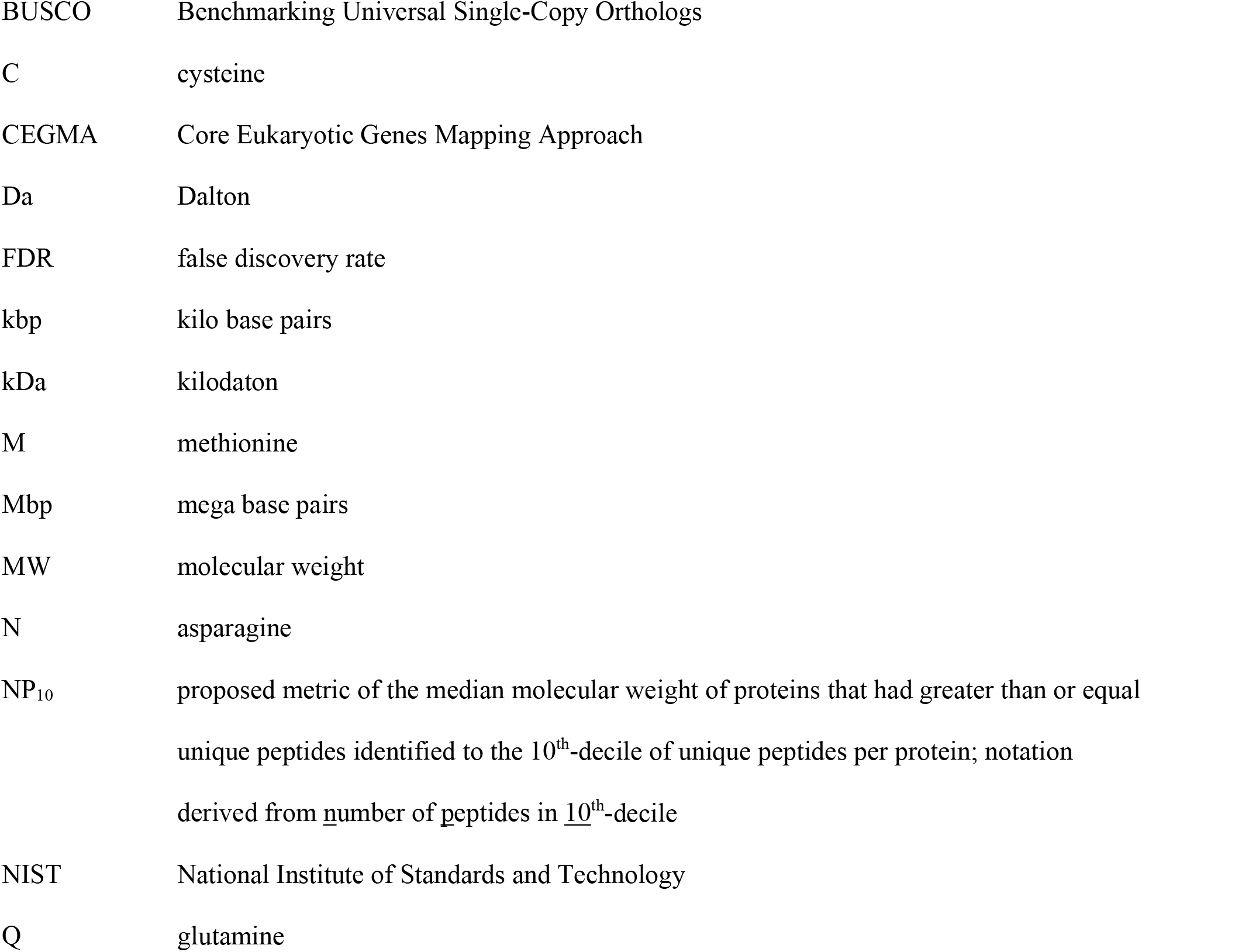

### Consent for publication

Not applicable.

### Authors’ contributions

All authors helped conceived of the study, developed methodology and assisted in reviewing the manuscript. DE and WD selected and processed samples for proteomic analysis and collected data. BN analyzed the data and wrote the initial manuscript draft.

## Acknowledgements

Specimens used for this study were collected by Wayne E. McFee (National Oceanic and Atmospheric Administration, National Ocean Service, National Centers for Coastal Ocean Science) and William A. McLellan (University of North Carolina, Wilmington) and provided by the National Marine Mammal Tissue Bank, which is maintained as part of the Marine Environmental Specimen Bank at NIST and is operated under the direction of NMFS and in collaboration with USGS, USFWS, BOEMRE (formerly MMS), and NIST through the Marine Mammal Health and Stranding Response Program. All samples were collected under approved permits issued to the MMHSRP (responsible party: Dr. Teri Rowles) and all sampling protocols were reviewed and approved by a NOAA/NMFS *ad hoc* Institutional Animal Care and Use Committee (IACUC). The authors wish to thank Michael G. Janech and staff at NIST for critical feedback.

## Disclaimer

Identification of certain commercial equipment, instruments, software or materials does not imply recommendation or endorsement by the National Institute of Standards and Technology, nor does it imply that the products identified are necessarily the best available for the purpose.

