## Supplementary figures and images for "Proteomics as a metrological tool to evaluate genome annotation accuracy following *de novo* genome assembly: a case study using the Atlantic bottlenose dolphin (*Tursiops truncatus*)"

### Supplementary Materials

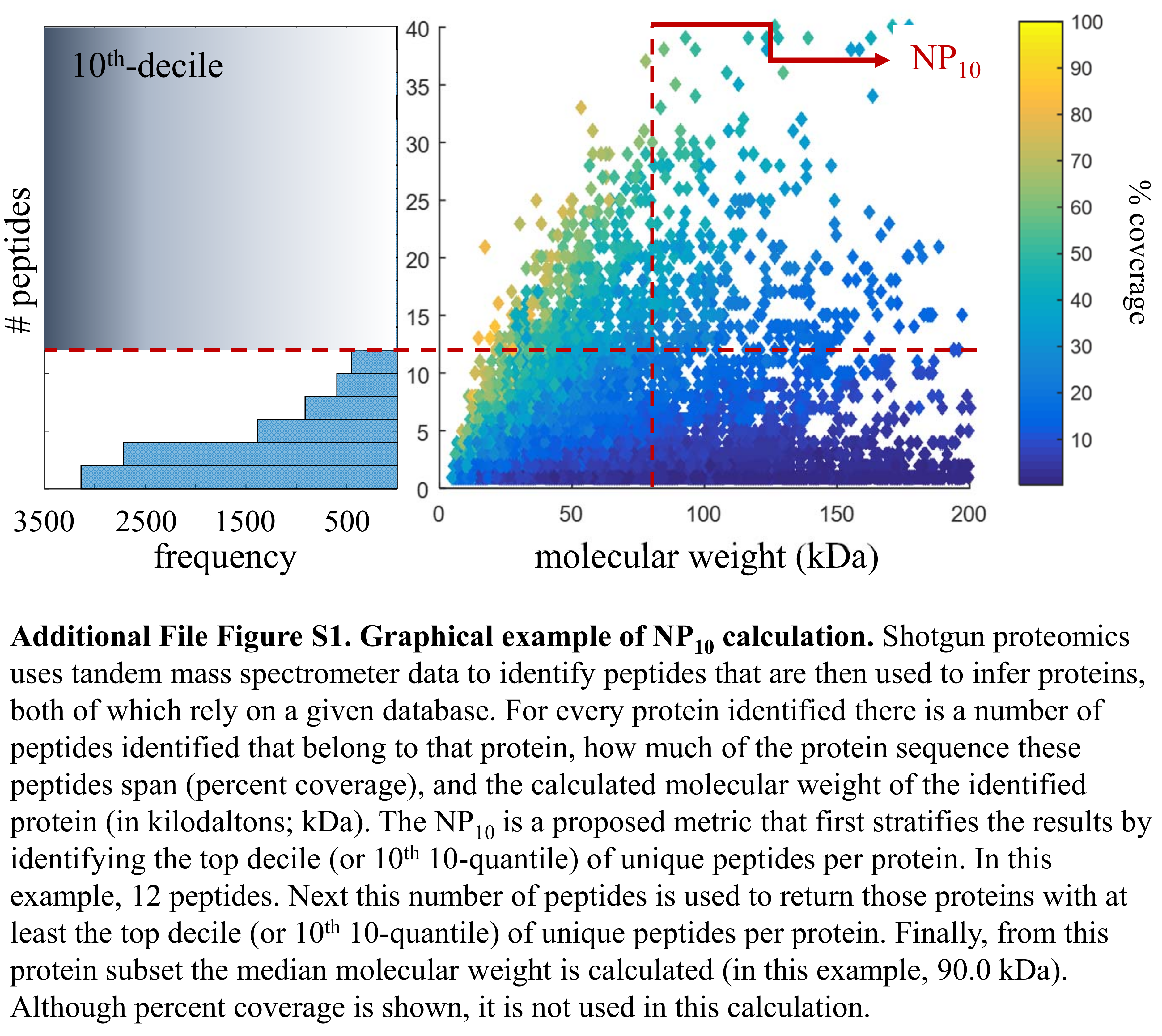
